# Reduced threat avoidance but increased stress induced approach bias in women taking oral contraceptives

**DOI:** 10.1101/2023.11.13.566589

**Authors:** Jasmin Thurley, Macià Buades-Rotger, Georg Serfling, Thessa Howaldt, Nicole Reisch, Ulrike M. Krämer

## Abstract

Recent research has increasingly acknowledged the impact of oral contraceptives on affective behavior and stress responses; however, the underlying mechanisms are still not well understood. Studies have previously shown that steroid hormones modulate automatic approach and avoidance behavior. Here, we thus investigated the effects of oral contraceptives on approach and avoidance behavior and whether these effects are modulated by stress. The study comprised 130 female participants, half of whom were using oral contraceptives, while the other half were not using any hormonal contraception (NC). The participants completed the Approach Avoidance Task (AAT), which measures automatic approach and avoidance behavior to socio-affective signals. The AAT was run once before and once after a stress manipulation using the Socially Evaluated Cold Pressor Test. OC users showed absent avoidance behavior to social threat signals and a stress-induced increase in approach behavior to positive social signals. The latter was found in particular in women taking androgenic acting OC, demonstrating that different OC preparations need to be taken into account in research on OC effects. However, OC and NC group did not differ in their cortisol stress response. Overall, the results suggest that OC usage impacts on approach and avoidance behavior to social signals, which might also contribute to the development of affective side effects.

## 1. Introduction

Oral contraceptives (OC) are amongst the most reliable methods to prevent undesired pregnancies and are the contraceptive of choice for many women worldwide (Christin-Maitre, 2013). Hormonal contraception, however, is associated with side effects, including changes in affective experience and a possibly increased risk for depression (Ernst et al., 2002; Gingnell et al., 2013; Montoya & Bos, 2017). Prevalence estimates of adverse effects on mood and affect are between 4 – 10% (Poromaa & Segebladh, 2012) with a higher risk among adolescents and women with previous mood-related side-effects of hormonal contraceptives (Lewis et al., 2019; Tronson & Schuh, 2022). Many OC users however also report improved mood and emotional well-being or no changes at all (Poromaa & Segebladh, 2012; Tronson & Schuh, 2022). The impact of oral contraceptives on specific neural and psychological mechanisms, which cause these changes in mood and affect, remains unclear. OCs can influence brain functions through multiple mechanisms: directly through binding of the synthetic estrogens and progestins to endocrine receptors, indirectly through a reduction of endogenous levels of estrogens and progesterone, and, finally, indirectly through effects on neurotransmitter systems or other endocrine receptors (androgen, glucocorticoid or mineralocorticoid receptors) (Tronson & Schuh, 2022).

Altered approach and avoidance behavior may be a possible psychological mechanism for several reasons: the brain regions which have been implicated in approach and avoidance behavior to socio-affective signals and its regulation, include especially the amygdala and anterior prefrontal cortex (Kaldewaij et al., 2017). OC users have been found to show functional and structural changes in these and neighboring brain regions (Brønnick et al., 2020; Gingnell et al., 2013; Petersen & Cahill, 2015; Sharma et al., 2020), which might contribute to altered approach/avoidance behavior. Converging evidence (Rehbein et al., 2021) suggests that amygdala activity to affective stimuli is modulated by estradiol (van Wingen et al., 2011; Zeidan et al., 2011), progesterone (van Wingen et al., 2007; van Wingen et al., 2008) and testosterone (Buades-Rotger et al., 2016; van Wingen et al., 2010). Moreover, the steroid hormones testosterone and cortisol, possibly modulated by OC use, appear to have an impact on approach and avoidance behaviors (Enter et al., 2014; Radke et al., 2015; Roelofs et al., 2005). Lastly, considering that OC influence the stress axis and cortisol reactivity (Mordecai et al., 2017), which in turn affects approach and avoidance behavior (Roelofs et al., 2005), it is possible that OC influence affective behavior in conjunction with stress. Here, we thus tested whether women who are taking OC show altered approach and avoidance behavior, and whether this effect is modulated by stress using a cross-sectional design.

Approach/avoidance behavior refers to the automatic tendency to approach positive stimuli and avoid negative stimuli, which can be measured using the Approach Avoidance Task (AAT) (Chen & Bargh, 1999). In one commonly used variant of the AAT (Kaldewaij et al., 2017), participants are asked to pull pictures of happy faces towards them and push negative faces away as fast as possible using a joystick. Response times in this congruent condition are compared with those of the reverse, incongruent condition (pull angry faces and push happy faces away) (Beyer et al., 2017). Typically, participants are faster in the congruent compared to the incongruent condition, referred to as congruency effect or AAT bias. Previous research with the AAT reported that people with social anxiety show faster avoidance (Roelofs et al., 2010), whereas psychopathic patients showed reduced avoidance to angry faces (von Borries et al., 2012), speaking for the validity of the AAT. A neurobiological model of the AAT (Volman et al., 2013) assumed that faces are first processed in the fusiform face area (FFA) and subsequently projected to the amygdala, which has a bidirectional connection to the anterior prefrontal cortex (aPFC). When automatic affective behavior needs to be controlled in the incongruent condition, amygdala activity is believed to be inhibited by the aPFC (Volman et al., 2013).

Oral contraceptive use has previously been found to be associated with changes in reactivity and activation of the PFC and amygdala. Women using OCs responded to negative emotional stimuli with reduced amygdala reactivity (Petersen & Cahill, 2015) and increased prefrontal activation (Sharma et al., 2020). Rodent studies suggest that estradiol treatment influences estrogen receptor density in the amygdala, whereby the direction of effects seems to depend on the amygdala nucleus and estrogen receptor subtype (ERα or ERβ) (Österlund et al., 1998). Estradiol has been shown to facilitate fear extinction learning through its effects in the amygdala and prefrontal cortex (Zeidan et al., 2011). Reduced circulating estradiol levels due to hormonal contraception in turn impaired extinction learning (Graham & Milad, 2013; Taxier et al., 2020). Although these findings cannot directly be translated to approach/avoidance behavior to social signals, they speak for reduced reactivity to threat signals in OC users. Moreover, Li and colleagues (Li et al., 2022) reported that in a sample of women not taking hormonal contraceptives, progesterone and estradiol levels were correlated with approach and avoidance behavior. Specifically, higher progesterone levels were related to faster avoidance and higher estradiol levels were related to faster approach behavior (Li et al., 2022). This would suggest reduced avoidance and approach behavior with the reduction of endogenous progesterone and estradiol in women taking OCs.

In addition to the contraceptive effect OC have on the HPG-axis (hypothalamic– pituitary–gonadal axis), they also affect the HPA stress axis (hypothalamic-pituitary-adrenal axis). One often replicated effect of OC use (Lewis et al., 2019; Tronson & Schuh, 2022) is a blunted cortisol stress response, which was first reported by Kirschbaum and colleagues (Kirschbaum et al., 1999; Kirschbaum et al., 1995). Additionally, several studies reported an increase of basal cortisol levels in OC users compared to women not taking hormonal contraceptives (Tronson & Schuh, 2022). This pattern of elevated basal cortisol levels together with a blunted cortisol stress response is also characteristic of depression as well as other psychiatric disorders (Carroll et al., 2017). However, in a large-scale study on OC-related changes in the stress axis, no association was found between basal plasma cortisol levels and depressive symptoms in women using OCs (Hertel et al., 2017), calling basal cortisol levels into question as a mediating factor between OC use and depression. Given the close interrelation between stress and motivation (Slattery & Cryan, 2017), it is however likely that a blunted stress response in OC users is associated with altered motivation and behavior. There is only few research on stress effects on the AAT, but this work suggests that a high cortisol increase due to social stress is associated with a reduced AAT congruency effect (Roelofs et al., 2005). As OC might also influence approach/avoidance behavior through this pathway, we tested whether approach/avoidance behavior in OC users interacts with stress exposure.

One challenge in research on OC is the large heterogeneity of preparations taken by women (Tronson & Schuh, 2022). In our study, only women taking combined preparations containing an estrogen and a progestin were included, but the preparations nevertheless differed in the included estradiol dose and in the used anti- or androgenic acting progestin. The different formulations could have an influence on AAT and stress effects via the different efficacy profiles, as there is for example evidence that women with androgenic OC have a more pronounced cardiovascular stress response (Straneva et al., 2000). Therefore, we performed exploratory analyses on the subgroups.

## 2. Methods

### 2.1 Participants

We tested one hundred thirty women, of whom sixty-five had been using oral contraceptives for at least three months (oral contraceptives, OC group) and 65 had not used any hormonal contraception for at least six months (no hormonal contraception, NC group). The mean age in the OC group was 21.9 years (SD = 2.13) and in the NC group 22.6 years (SD = 2.4). They had all normal or corrected to normal vision (self-report). An analysis with G*Power 3.1.9.6 (Faul et al., 2007) suggested that at least 128 participants would be required to find an effect of medium effect size for group differences in the AAT effect (f = 0.25) with a power of 0.8 and an alpha error of 0.05. All participants were between 19 and 30 years old, had a BMI between 17.5 and 25 and no neurological, psychiatric, cardiovascular or endocrinological pre-existing diseases.

Of the 130 participants, we had to exclude three for the following reasons: one due to many errors in the AAT (mean error rate: 39.15%) and two due to reaction times in the AAT that were more than 3 SD above the mean. Saliva samples from 20 participants could not be used, resulting in a final sample of 106 participants (50=OC, 56=NC) for the cortisol analyses.

All of the women in the OC group were using combined oral contraceptives containing both estradiol and progestin. The doses of ethinyl estradiol varied between 20 and 30 µg. Three women were using OC with estetrol (14.2 mg) as estradiol and 4 women took OC containing estradiol valerate (1-3 mg). The doses of progestin ranged between 100 and 3000 µg (see Supplementary Table 1). The participants were recruited via mailing lists of the University of Lübeck. For the verification of the inclusion and exclusion criteria, the potential participants were subsequently called. The person who conducted the calls was not involved in the measurements and the investigators were blinded to the group membership of the respective participants. All participants were informed about the study objectives and compensated for their participation with 10 Euros per hour or credit points. All participants gave their written informed consent in accordance with the Declaration of Helsinki. The Ethics Committee of the University of Lübeck approved this study protocol (22-044).

### 2.2 Design and Procedure

OC women were tested between the first and third day of pill intake, NC women correspondingly on the eighth to tenth day of the menstrual cycle (follicular phase). The timing was chosen because interpersonal variations in cycle length (in the NC group) and associated differences in hormone concentrations are lowest at the beginning of the cycle (Hampson, 2020) and to measure women taking OC already during the active phase of pill intake. All measurements took place in the afternoon between 2 and 7 p.m., since during this period salivary cortisol concentrations are relatively consistently low for both women taking oral contraceptives and women with no hormonal contraception (Meulenberg & Hofman, 1990).

After participants gave their informed consent, they were asked to fill out some computerized questionnaires assessing emotion regulation, personality, and approach/avoidance tendencies (see 2.6 Questionnaires and Demographic Data). In the next step, they provided a saliva sample (t_0_) before performing the AAT on the computer (Figure 1). After another saliva sample (t_1_), the Socially evaluated cold pressor test (SECPT) (Schwabe & Schächinger, 2018) was performed. After a break of approximately 12 minutes, the AAT was performed a second time. After completion of the second AAT, another saliva sample (t_2_) was collected and the participants filled in a questionnaire with demographic data and information about the pill. Lastly, participants were debriefed about the research question and were compensated. The testing lasted approximately one hour in total.

**Figure 1.**
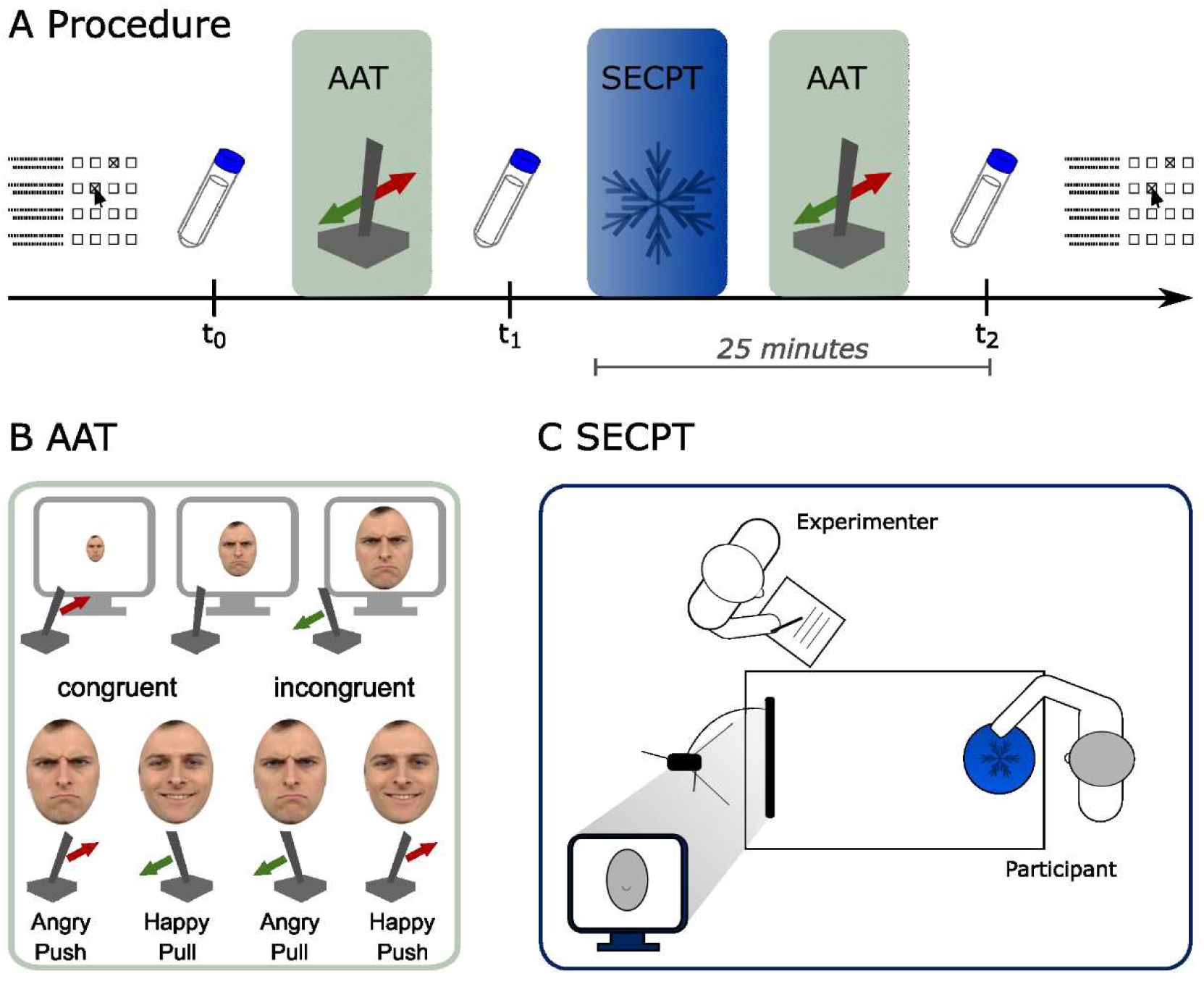
**A** Procedure. At first, the participants completed the questionnaires ERQ, BFI-10, and BIS/BAS. Next, they provided a saliva sample and performed the AAT (Approach Avoidance Task). After another saliva sample, they performed the Socially Evaluated Cold Pressor Test (SECPT) and another run of the AAT. 25 minutes after the onset of the SECPT, the participants gave another saliva sample and finally provided demographic information. **B** Approach Avoidance Task (AAT): Participants were asked to push away angry faces and pull happy faces as fast as possible in the congruent condition and do it the other way round in the incongruent condition. Note that drawings are shown for visualization only. In the experiment, we showed images of happy and angry faces, which were taken from the Radboud Faces database (Langner et al., 2010). **C** Socially evaluated cold pressor test (SECPT). Participants were required to keep their dominant hand in ice water for 3 minutes while being observed by the experimenter and being recorded on video.

### 2.3 Approach Avoidance Task (AAT)

We used the Approach Avoidance Task (Chen & Bargh, 1999) in the version of Beyer et al. (2017) with images of happy and angry faces from the Radboud Faces database (Langner et al., 2010) as stimuli. Stimuli were presented using Presentation® (Neurobehavioral Systems). In total, images of thirty individuals (15 male, 15 female) were shown during the AAT. There was one picture of each person with an angry and a happy facial expression. For the practice blocks, pictures of nine different people than in the main block were used. All pictures were cut into an oval shape so that hair, neck and ears are not visible.

Participants used the joystick with their dominant hand. Before each stimulus, a small black cross appeared on the white screen. As soon as the “shoot” button of the joystick was pressed, the cross disappeared and a face appeared in the center of the screen. By pushing the joystick away (avoidance response), the image got smaller step-by-step (seven steps in total). Pulling the joystick towards the body (approach response) made the image larger. Two conditions were run in two different blocks. In the congruent condition, participants were asked to pull happy faces toward them as quickly as possible and to push angry faces away from them. In the incongruent condition, the instruction was reversed. Block order was kept identical across participants and stress conditions, starting always with the congruent condition. Here, happy faces were to be pushed away from the body and angry ones were to be pulled in. Each block contained thirty happy faces and thirty angry faces, using the same images in both blocks.

Before each block, the participants could familiarize themselves with the task in a practice session. Here, the participants received direct feedback whether they had reacted correctly (green tick for correct reaction and red cross for incorrect reaction). The first exercise run consisted of twenty images. In the second practice run, 28 pictures were used because the instruction that was internalized in the first block was reversed and task-switching effects were to be avoided.

During each trial the reaction time was measured, which is determined as the interval between the appearance of the stimulus and the start of the movement of the joystick.

### 2.4 Socially Evaluated Cold Pressor Test (SECPT)

The SECPT was performed as suggested by Schwabe and Schächinger (2018). The participants were instructed to hold their dominant hand, including the wrist, in a bucket of ice water (0-2 degrees Celsius) until the experimenter asked them to take the hand out again (they were not told in advance how long they should keep the hand in the ice water). Only if they could no longer stand the cold at all, they were allowed to take their hand out of the water earlier. They were also told that they were videotaped during the experiment so that their facial expressions can be analyzed later. Therefore, they were instructed to look into the camera during the entire experiment and not to speak. The camera was placed about two meters in front of the participants behind a screen facing them. The camera recorded the subject’s face and transferred it directly to the screen so that the participant could see herself. Next to the camera, the experimenter stood at a distance that allowed the participant to simultaneously look into the camera and see the experimenter out of the corner of her eye. The experimenter made sure that the participant did not make a fist and that her hand was in ice water up to the wrist. At the same time, she took notes and avoided giving confirming signals such as smiling or nodding. After three minutes, the test ended and the participant was allowed to take her hand out of the water. If she took her hand out before that, the experimenter responded by asking the participant to put her hand back in the water. If this was not possible, the participant was asked to remain standing and looking into the camera for the remainder of the three minutes.

### 2.5 Saliva Sampling

Saliva samples were collected at three time points of the measurement. For this purpose, we used Sarstedt cortisol Salivette® with a synthetic fiber roll. To prevent cortisol levels from being influenced by factors other than the experimental factors, participants were instructed not to exercise, eat anything, drink coffee, take medications, or smoke for one hour before the appointment (Roelofs et al., 2005). Saliva samples were frozen and stored at -20°C until analysis. After thawing, samples were centrifuged at 3,000 rpm for 5 min, which resulted in a clear supernatant of low viscosity. Salivary concentrations were measured using commercially available chemiluminescence immunoassay with high sensitivity (Tecan - IBL International, Hamburg, Germany; catalogue number R62111). The intra and interassay coefficients of variance were below 9%. Samples were considered empty if there was no saliva in the tube after centrifugation (samples from n=20 participants, resulting in a sample of 106 participants (50=OC, 56=NC) for the cortisol analyses).

### 2.6 Questionnaires and Demographic Data

We used the Behavioral Inhibition Scale (BIS) and the Behavioral Approach Scale (BAS) designed by Carver and White (1994) to assess tendencies to exhibit avoidance or approach behaviors. We also used the Emotion Regulation Questionnaire (ERQ) (Gross & John, 2003) and the Big Five Inventory (BFI-10) (Rammstedt & John, 2007) to examine interpersonal differences in emotion regulation and personality structure.

Participants also provided information on the name of the OC preparation taken, previous duration of use and age at first use of oral contraceptives, time of daily use, reason for use, and any side effects (OC group). A summary of the frequency of reported side effects in our sample is provided in Supplementary Table 2. The NC-women were asked if they had used hormonal contraception in the past. If they had taken OC in the past, the duration and the time since the last intake was recorded. The reason for taking the pill and the occurrence of any side effects were asked as well. The weight, height and age of all test participants was recorded.

### 2.7 Data analysis

The main outcome of the AAT were the reaction times (RT) which were defined as the interval between stimulus presentation and movement onset. Following previous work (Beyer et al., 2017), we excluded trials with response latencies of less than 150 ms or more than 3 SD from the individual participant’s own mean. Moreover, we only considered correct trials. Incorrect trials (errors), which also included trials with an initial movement into the wrong direction and a following correction were excluded. We first log-transformed the reaction times (ms) before computing mean response times for each condition (push, pull for angry and happy) and participant, as reaction times were not normally distributed. We also considered the error rates for determination of outliers. Participants (total n=3) were excluded if reaction times (n=2) or error rates (n=1) in the AAT were more than 3 SD above the mean across all participants.

We computed RT means, outliers and error rates using MATLAB (MATLAB, 2021). Statistical analyses were conducted in Jamovi (The jamovi project, 2021) and JASP (JASP Team, 2023).

We performed a mixed effects ANOVA that included the factors emotion (happy, angry), movement (push, pull), stress (within-subject factors) and contraception (between-subject factor) using log-transformed reaction times. To parse interaction effects, we conducted further ANOVAs to examine group differences between OC and NC women separately for happy and angry faces and before and after stress. To determine the AAT effect, the reaction times of the movement away from the body (push) were subtracted from the reaction times of the movement towards the body (pull). A higher (more positive) score here reflects stronger avoidance behavior and a lower (more negative) score reflects approach behavior. The AAT effect was calculated as the difference between the bias score for angry faces and the bias score for happy faces. Accordingly, the higher the bias scores, the larger the effect. The AAT effect was reported only for visualization and to allow for comparability of results with previous studies and is not interpreted on its own in this study. We conducted an additional mixed effects ANOVA with factors stress and contraception to compare the error rates of OC and NC women before and after stress.

To investigate the change in cortisol levels, we performed another mixed effects ANOVA. Here, the within-subject factor time (t_0_, t_1_, t_2_) and the between-subject factor contraception were included. For the comparison of group differences in the questionnaires, we calculated two-sided independent samples t-tests.

For further exploratory analyses of subgroups within the OC group, we ran additional mixed effects ANOVAs with the factors emotion, movement, stress (within-subject factors) and the respective between-subject factors (androgenicity, EE-dose, onset, time of OC intake or contraception). Moreover, we ran a correlation of duration of OC intake and the AAT effect and a correlation of the AAT effect with the Emotion Regulation Questionnaire subscales (across the whole sample).

All results with a p-value less than .05 were considered significant. For significant ANOVA results, we used Bonferroni-Holm adjusted p-values for post-hoc tests. In order to make informed statements regarding the evidence, we also provide the Bayes Factor, which tells us how much more probable the observations are under the alternative hypothesis compared to the null hypothesis. Our Bayesian analysis was conducted using the default priors in JASP, and we present the findings of Bayesian model averaging (BF_incl_) (Love et al., 2019). To interpret the results, we adhere to established guidelines: BF values exceeding 3 indicate moderate support, while values above 10 indicate strong support for the alternative hypothesis. Conversely, BF values below 0.1 strongly support the null hypothesis, and values below 0.33 suggest moderate support (van Doorn et al., 2021). Because Bayesian analyses are time-consuming and computationally expensive for large-sample multifactorial ANOVAs, we report Bayes factors exclusively for specific lower-order effects.

Data is available at https://osf.io/kz4yp/?view_only=167cd3c80d064916ae0edcd82f02b5ac.

## 3. Results

### 3.1 Approach Avoidance Behavior

The AAT showed the expected experimental effects, as participants showed avoidance of angry faces and approach toward happy faces. Visual examination of the behavioral data suggested, however, that OC women responded less avoidant overall and even showed a slight tendency to approach angry faces before the SECPT (see Table 1 and Figure 2A&B). Note that the reaction times were log-transformed to increase normality.

**Table 1.** Sample descriptives

| | NC ($n=65$) | OC ($n=65$) | $t(128)$ | $p$ | $d$ | BF<sub>10</sub> |
| --- | --- | --- | --- | --- | --- | --- |
| Age | 22.6 ± 2.40 | 21.9 ± 2.13 | 1.58 | .116 | .278 | 0.583 |
| BMI | 21.3 ± 1.72 | 21.1 ± 1.77 | 0.809 | .420 | .143 | 0.253 |
| *Questionnaires* |  |  |  |  |  |  |
| BIS | 2.09 ± 0.063 | 2.06 ± 0.055 | 0.368 | .713 | .065 | 0.199 |
| BAS | 1.87 ± 0.039 | 1.91 ± 0.038 | -0.836 | .405 | -.147 | 0.258 |
| BFI-10 (openness) | 3.76 ± 0.125 | 3.59 ± 0.123 | 0.965 | .336 | .169 | 0.286 |
| BFI-10 (neuroticism) | 3.22 ± 0.110 | 3.45 ± 0.114 | -1.460 | .148 | -.255 | 0.490 |
| BFI-10 (extraversion) | 3.62 ± 0.112 | 3.62 ± 0.118 | 0.047 | .962 | .008 | 0.188 |
| BFI-10 (conscientiousness) | 3.89 ± 0.100 | 3.77 ± 0.094 | 0.897 | .371 | .157 | 0.270 |
| BFI-10 (agreeableness) | 3.71 ± 0.109 | 3.49 ± 0.113 | 1.371 | .173 | .240 | 0.440 |
| ERQ (cognitive reappraisal) | 4.91 ± 0.122 | 4.66 ± 0.107 | 1.522 | .130 | .267 | 0.535 |
| ERQ (expressive suppression) | 3.17 ± 0.149 | 3.24 ± 0.133 | -0.347 | .729 | -.061 | 0.198 |

Notes: BIS Behavioral Inhibition Scale, BAS Behavioral Approach Scale, BFI Big Five Inventory, ERQ Emotion Regulation Questionnaire; values are mean ± standard deviation

**Figure 2.**
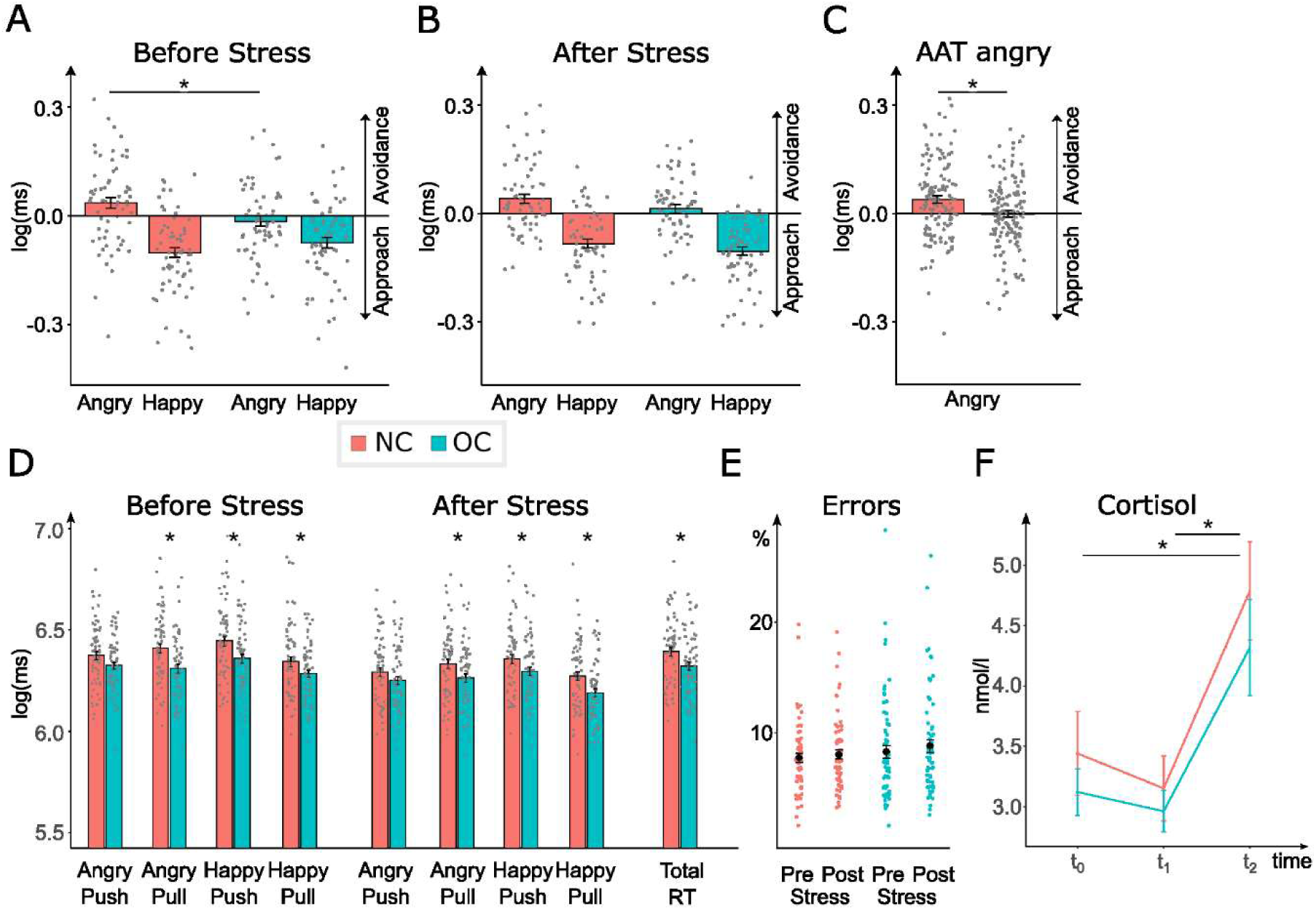
**(A/B)** Approach Avoidance Task (AAT) - Bias Scores (log-transformed) for angry and happy faces for women with no hormonal contraception (NC; red) and women taking oral contraceptives (OC; blue) before stress induction through Socially Evaluated Cold Pressor Test (SECPT) **(A)** and after stress exposure **(B). (C)** AAT – Bias Scores (log-transformed) for angry faces across both AAT-runs for NC (red) and OC (blue) women. **(D)** AAT - Reaction times (log-transformed) for each emotion (happy or angry), movement (push or pull) and mean reaction times over all trials before and after stress for NC and OC women. **(E)** Error rates of runs of the AAT for each participant in percent before and after stress. **(F)** Cortisol levels (log-transformed) at three measuring points for NC and OC women. Error bars indicate standard error, asterisks indicate significant differences (p < 0.05).

Results of the mixed ANOVA showed that, as expected, the responses to the emotions in the congruent and incongruent condition differed in speed (see Figure 2A-D). Overall, it took the participants longer to pull angry faces towards them than to push them away, whereas they showed the opposite pattern for happy faces (emotion x movement: *F*(1,125) = 66.733, *p* < .001, *η^2^_p_* = 0.348), reflecting the expected AAT effect. In general, the groups did not differ in the expression of this pattern (emotion x movement x contraception: *F*(1,125) = 2.482, *p* = .118, *η^2^* = 0.019). However, there was a significant fourfold interaction of stress x emotion x movement x contraception (*F*(1,125) = 4.345, *p* = .039, *η^2^* = 0.034). Similar to previous studies with the AAT (Radke et al., 2015; Volman et al., 2013), participants were faster in pull than push movements (movement: *F*(1,125) = 95.735, *p* < .001, *η^2^* = 0.434). Interestingly, OC women responded overall faster than NC women (contraception: *F*(1,125) = 6.35, *p* = .013, *η^2^* = 0.048) (see Figure 2D). Women in the OC group were especially faster than NC women in their pull movements, and less so in their push movements (interaction of movement x contraception: *F*(1,125) = 6.019, *p* = .016, *η^2^* = 0.046; pull: *t*(125) = 2.827, *p*_Holm_ = .016, *d* = 0.461; push: *t*(125) = 2.166, *p_Holm_* = .064, *d* = 0.353).

To understand the four-way interaction of stress x emotion x movement x contraception, we computed separate mixed effects ANOVAs for the two emotions (happy and angry), since we expected differences especially for angry faces. In response to angry faces, the groups indeed differed in their behavior, reflected in a significant interaction of movement x contraception (*F*(1,125) = 6.708, *p* = .011, *η^2^* = 0.051, *BF* = 3.916). The Bayes factor indicated moderate support for this interaction. While NC women pushed away angry faces faster than they pulled them closer (p_Holm_ = 0.003), showing the typical avoidance bias, OC women responded equally fast when pushing and pulling (p_Holm_ = 0.937; see Figure 2C). This pattern was not influenced by stress (stress x movement x contraception: *F*(1,125) = 1.155, p = .285, *η^2^* = 0.009, *BF* = 0.305). The groups did not differ in their general behavior to happy faces (movement x contraception: *F*(1,125) = 0.039, *p* = .843, *η^2^* < 0.001, *BF* = 0.216), however, there was a threefold interaction of stress x movement x contraception for happy faces (*F*(1,125) = 6.596, *p* = .011, *η^2^* = 0.050, *BF* = 3.073). To understand this interaction, we computed separate mixed effects ANOVAs for the two groups (NC and OC). While stress had no influence on approach behavior to happy faces in NC women (stress x movement: *F*(1,62) = 1.56, *p* = .216, *η^2^* = 0.025, *BF* = 1.419), OC women differed in their reactions towards happy faces before and after stress (stress x movement: *F*(1,63) = 5.68, *p* = .02, *η^2^* = 0.083, *BF* = 9.59). This was due to more pronounced approach behavior after stress than before stress (movement after stress: *F*(1,63) = 87.2, *p* < .001, *η^2^* = 0.580, *BF* = 6069.494; movement before stress: *F*(1,63) = 27.1, *p* < .001, *η^2^* = 0.301, *BF* = 2.379×10^10^) (see Figure 2A&B). Note however, that OC women showed a clear approach bias to happy faces both before and after stress.

Regarding error rates in the AAT, there was no group difference before or after stress (mixed effects ANOVA with factors stress and contraception; contraception: *F*(1,125) = 1.05, *p* = 0.308, *η^2^* = 0.008, *BF* = 0.257) (Figure 2E).

### 3.2 Stress effects on cortisol level

The stress manipulation generally succeeded, as cortisol levels were higher on average after the SECPT than before. About 13% of the participants discontinued the SECPT early, which is comparable to the data presented by Schwabe and Schächinger (2018). Since those participants in previous studies did not differ in their cortisol response from those who persisted for the three minutes in the stress situation (Schwabe & Schächinger, 2018), they were not excluded from further analyses.

Cortisol values were log-transformed to obtain a normal distribution. Additionally, we performed a Greenhouse Geisser correction due to the violation of the assumption of sphericity. Cortisol levels differed significantly at the three measurement time points (two before SECPT and one after) (time: *F*(2,214) = 41.433, *p* < .001, *η^2^* = 0.279, *BF* = 1.076×10^13^). As expected, we observed a strong increase in cortisol after the stress manipulation (t_2_) compared to directly before (t_1_) (*t*(107) = -8.452, *p_Holm_* < .001, *d* = -0.647, BF = 3.983×10^8^) and compared to the first measurement (t_0_) (*t*(107) = -7.154, *p_Holm_* < .001, *d* = -0.548, BF = 256993.046) . However, there was no significant difference between the NC and OC women in cortisol levels (contraception: *F*(1,107) = 0.201, *p* = .655, *η^2^* = 0.002, *BF* = 0.231) and no significant interaction of time and contraception (*F*(2,214) = 0.547, *p* = .580, *η^2^* = 0.005, *BF* = 0.108) (see Figure 2F). We also computed the area-under-the-curve values AUC-G and AUC-I as suggested by Pruessner and colleagues (Pruessner et al., 2003), but these values did not differ between the groups either (both t < 1 and p > 0.5).

### 3.3 Exploratory Analyses

#### Composition of preparations

OC preparations differed in composition with regard to the type of progestin and the dose of ethinyl estradiol (EE) they contain. The partial effects of the progestin can be distinguished into androgenic and anti-androgenic effects. To compare OC women with androgenic (N=27, mean(SD) progestin dose: 122(100)) and anti-androgenic (N=37, mean(SD) progestin dose: 2108 (315)) preparations, we conducted a mixed effects ANOVA (within factor: stress, emotion, movement; between factor: androgenicity). The results showed a significant four-way interaction of stress x emotion x movement x androgenicity (*F*(1,62) = 6.380, *p* = .014, *η^2^* = 0.093). Separate ANOVAs for the emotions happy and angry showed that only for happy faces there was a significant three-way interaction of movement x stress x androgenicity (*F*(1,26) = 6.502, *p* = .013, *η^2^* = 0.095, *BF* = 1.056). Using further separate ANOVAs for androgenic and anti-androgenic groups with factors stress and movement, we observed that only for women taking androgenic acting OC there was a significant interaction of stress and movement (*F*(1,26) = 19.162, *p* < .001, *η^2^* = 0424, *BF* = 269.14). Post hoc tests showed that women taking androgenic acting OC had a stronger approach bias for happy faces after stress than before (before stress: (*t*(26) = 2.726, *p_Holm_* = 0.022, *d* = 0.371, *BF =* 2.501); after stress: (*t*(26) = 6.199, *p_Holm_* < 0.001, *d* = 0.843, *BF =* 96212.845)). This suggests that the stress effect on the approach bias in OC reported above was due in particular to the behavior in the group of women taking androgenic acting OCs. The overall response times, however, did not differ between the androgenic and anti-androgenic groups (*F*(1,62) = 1.126, *p* = .293, *η^2^* = 0.018, *BF* = 0.179) (Figure 3A&B).

**Figure 3.**
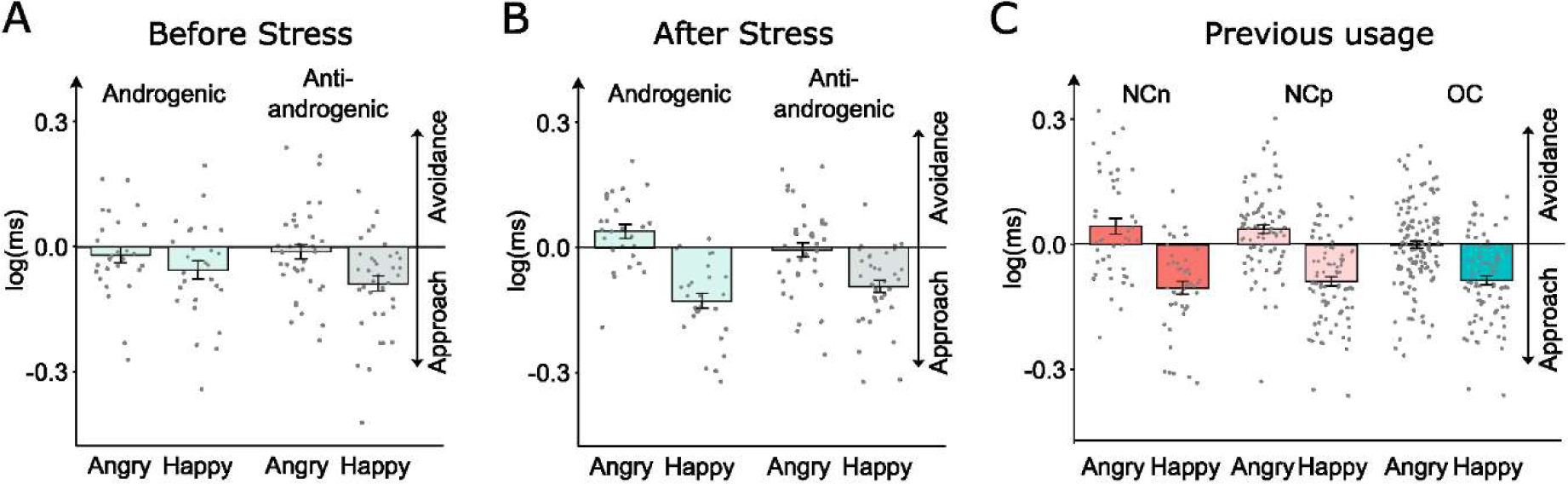
**(A/B)** Approach Avoidance Task (AAT) - Bias Scores (log-transformed) for angry and happy faces for women taking oral contraceptives (OC) with androgenic progestin (light green) and those with anti-androgenic progestin (light grey) before stress induction through Socially Evaluated Cold Pressor Test (SECPT) **(A)** and after stress exposure **(B)**. **(C)** Bias Scores (log-transformed) for angry and happy faces for women who never took OC (NCn, red), who took OC previously but discontinued (NCp, pink) and OC women (blue) over both AAT-trials. Error bars indicate standard error.

Most OC preparations used in our sample contained 30µg (EE30) (N=41, one excluded in analysis) or 20µg (EE20) (N=17) of ethinyl estradiol. Note that the ethinyl estradiol dosage was not independent of the progestin type: all EE20 preparations were combined with an androgenic acting progestin. The EE30 preparations were combined with either androgenic (N=10) and anti-androgenic (N=30) acting progestins. When conducting a mixed effects ANOVA (within factors: stress, emotion, movement; between factor: EE-dose), we again found a fourfold interaction of stress x emotion x movement x EE-dose (*F*(1,55) = 4.267, *p* = .044, *η^2^* = 0.072). Again, this reflected a stronger approach bias to happy faces after stress in women with a low EE dosage (and androgenic preparation) (interaction of stress and movement: *F*(1,16) = 14.52, *p* = .002, *η^2^_p_* = 0.476, *BF* = 56.229; movement effect before stress: *t*(16) = 1.0, *p_Holm_* = 0.399, *d* = 0.207, *BF =* 0.347; movement effect after stress: *t*(16) = 4.213, *p_Holm_* < 0.001, *d* = 0.871, *BF =* 262.054).

The women did not differ in their cortisol response as a function of either androgenicity or EE dose (no significant interaction of time and androgenicity/ EE dose: F(2,98) = 0.566, p = 0.596, *η^2^* = 0.011 / F(2,88) = 0.434, p = 0.649, *η^2^* = 0.01).

#### Conditions of OC intake

Previous research suggests that certain conditions of OC use, such as age at first use or whether OC were taken before or after the measurement, may be important (Anderl et al., 2022; Gravelsins et al., 2021). In our sample, participants did not differ in their approach and avoidance behaviors depending on whether they started taking OC as adolescents (aged younger than 18 years (N=42)) or as adults (aged older than 18 years (N=22)) (no interaction of stress x emotion x movement x onset: *F*(1,62) = 0.997, *p* = .322, *η^2^* = 0.016), nor did the timing of daily intake (before or after measurement) make a difference (no interaction of stress x emotion x movement x time of OC intake: *F*(1,62) = 2.6674, *p* = .107, *η^2^* = 0.041). We were also interested in whether the duration of intake correlated with the expression of the AAT effects, which was not the case (*r* = -0.047, *p* =.710, *Fisheŕs z* = 0.128, *BF* = 0.167).

#### Previous OC intake

The work of Pletzer and Kerschbaum (2014) suggested that psychological effects of OC use may not be fully reversible. As some women in the NC group had previously taken oral contraceptives (N=41), we tested this for our data using another mixed effects ANOVA (within factors: stress, emotion, movement; between factor: contraception). The factor contraception had three levels: OC, NC - previously taken OC (NCp, N=41) and NC who had never taken OC (NCn; N=22). We found a significant main effect of contraception on response times (*F*(2,124) = 4.954, *p* = .009, *η^2^* = 0.074). The Post Hoc analysis shows that only the NCn women were slower than OC women in the AAT (*t*(124) = 3.117, *p* = .007, *d* = 0.699), while NCp and OC women did not differ (*t*(124) = 1.402, *p* = .164, *d* = 0.255). However, there was no significant interaction of emotion x movement x contraception (*F*(2,124) = 1.397, p = .251, *η*^2^*_p_* = 0.022)) (see Figure 3C).

### 3.4 Questionnaires

The groups did not differ significantly from each other in their scores in the BIS or BAS, BFI-10, or ERQ scales (see Table 1), suggesting that there were no general personality differences between NC women and OC users. The ERQ and the BIS and BAS subscales did not correlate with the AAT bias scores (all p > 0,05), neither did the BFI-10 subscales.

## 4. Discussion

Oral contraceptives have an effect on central nervous levels of steroid hormones, which in turn are known to modulate affective behavior. In the current study, we investigated whether OC usage is related to altered approach and avoidance behavior in a between-subject design. The results of this study suggest that women using oral contraceptives (OC) have significantly reduced avoidance tendencies to social threat signals compared to women not using oral contraceptives (NC). OC women were as quick to pull angry faces towards them as they were to push them away and thus showed no avoidance tendencies. Socially evaluated physical stress led to a similar rise in cortisol levels in both groups, but only in the OC group did stress increase approach behavior to positive social signals. It was also noteworthy that women taking OC responded overall faster than the NC women. Additional exploratory analyses suggested that the composition of the preparations may have influenced the AAT results. Only the OC women with an androgenic preparation showed a significant stress effect on approach behavior. The composition of preparations was not related to the decreased avoidance effect to threat signals in OC users, however. In sum, our study adds to the existing literature on how contraceptives can impact affect in women, and points to altered approach/avoidance tendencies as potential underlying mechanism of changes in affect and behavior.

### 4.1 OC-related differences in approach and avoidance behavior

We found that OC women pulled angry faces significantly faster towards them than NC women did, but were comparably fast at pushing them away. This suggests that OC primarily facilitate approach toward threat signals. Additionally, OC women responded with stronger approach tendencies to happy faces after stress than before. These differences were found relative to NC women who were in their early to mid follicular phase.

These results expand on recent studies on emotional reactivity changes in relation to OC usage (Lewis et al., 2019; Montoya & Bos, 2017). For instance, Hamstra et al. (2014) reported that women taking oral contraceptives detected fewer facial expressions of anger than NC women, but did not differ in recognizing happy faces. Although we did not find a reduced error rate, OC women did show a reduced avoidance response to angry faces. This might be related to the previously reported OC-related reduction in amygdala activation to negative stimuli (Petersen & Cahill, 2015) and reduced ERP responses to aversive stimuli (Monciunskaite et al., 2019), but as we did not measure neural activity, this remains speculative. Our results are in contrast to other studies, suggesting *enhanced* emotional reactivity in OC users, especially to negative stimuli (Radke & Derntl, 2016; Spalek et al., 2019). However, these studies measured explicit evaluations of the emotional stimuli, whereas the AAT aims to measure automatic behavioral tendencies. Crucially, our data thus point to motivational implications of OC usage, leading to altered affective action control and not mere perceptual differences. Future studies should combine behavioral measures of automatic affective behavior with neural measurements to tap into the underlying neural networks which are affected by OCs.

The AAT effects we observed were independent of the exact OC preparation. As we did not select for specific OC preparations and did not measure hormonal levels, conclusions regarding the relative role of estradiol and progesterone are difficult. Previous research has suggested that endogenous estradiol is associated with increased activity in emotion-regulatory brain areas such as the dorsolateral prefrontal cortex or the anterior cingulate cortex (Chung et al., 2019; Sharma et al., 2021). For progesterone it is known that it impairs emotion processing and recognition (Derntl et al., 2008; Guapo et al., 2009; van Wingen et al., 2007). Based on these previous findings, the observed AAT effects might rather be attributed to the OC effects on estradiol levels, but this needs to be investigated in future studies.

OC women were also generally faster than NC women. This is similar to results of another study, who also reported faster response times in OC users, this time to visual stimuli without any emotional component (Grikšienė & Rukšėnas, 2009). Older studies, however, reported generally increased reaction times in OC users (Garrett & Elder, 1984; Wuttke et al., 1975). However, it is important to note that other formulations of oral contraceptives were used at the time the studies were conducted. In a more recent study presenting emotional stimuli, women taking OC were also faster than NC women, but only in response to sad faces and not for happy or angry faces (Hamstra et al., 2014). Moreover, there is some evidence that the menstrual cycle phase has an influence on the reaction times in various paradigms (Li et al., 2022; Pletzer et al., 2014). Group differences in response times independently of experimental conditions are notoriously difficult to interpret as various factors might have contributed, such as arousal, psychomotor processing speed or motivation. It remains for future studies to examine how reliable these response time differences are.

In our study, women who had previously taken OC (NC_p_) did not differ in their approach avoidance pattern from women who had never taken OC (NC_n_). Regarding general response times however, women who had never taken OC were slowest, whereas women who had taken OC previously were not slower than current OC users. Pletzer and Kerschbaum (2015) suggested that effects of oral contraceptives could persist beyond the duration of intake. While they observed neurostructural changes after OC use (relative hippocampal gray matter volume was found to be associated positively with the duration of previous OC use), we found an effect on a behavioral level. Although the effect was of medium size, it should be treated with caution because of the unequal group sizes and the exploratory nature of the analysis. Nevertheless, the results show that, in future studies, a differentiation between current and previous OC users and nonusers might be useful.

### 4.2 OC-related effects in their interaction with stress

As hypothesized, stress influenced approach/avoidance behavior of OC and NC women in different ways. Only the OC women showed altered AAT behavior after stress compared to before. They reacted after stress with stronger approach tendencies to happy faces and unaltered to angry faces. The increased approach bias resulted primarily from faster pull reactions to happy faces. Our hypothesis was based on previous work by Roelofs and colleagues (2005), who showed reduced approach and avoidance biases after stress in high cortisol responders. Our results are only partly in line with these findings as the stress effects in our sample were independent of the cortisol response and specific for approach behavior. To our knowledge, no other study tested for stress effects on approach or avoidance biases in women taking oral contraceptives. Our results thus remain to be replicated.

The stress induced increase in the approach bias in OC users could be interpreted as kind a of tend-and-befriend response, characterized by responding to stress by aligning with social groups to ensure security (Taylor et al., 2000). Although the tend-and-befriend theory assumes that steroid hormones together with oxytocin drive the behavioral response to stress (Taylor et al., 2000), there is to our knowledge little direct evidence for estradiol or progesterone influencing social behavior after stress. In fact, one study revealed no significant difference between OC and NC women in prosocial behavior under stress (von Dawans et al., 2019). However, the study did not distinguish between OC women using androgenic and anti-androgenic preparations. In our study, we noticed that women taking androgenic OC reacted with heightened approach to happy faces after stress. There is some evidence that women using androgenic OC showed increased stress effects on vasoconstriction compared to women using anti-androgenic OC (Straneva et al., 2000). However, so far there is no study directly comparing androgenic and anti-androgenic OC effects on cognitive and affective changes after stress.

Indirect evidence for a role of steroid hormones on stress-induced changes in social cognition and behavior comes from studies showing sex differences in the response to stress. Women showed improved affective self-other distinction after stress (Tomova et al., 2014), whereas men showed worse self-other distinction resulting in an increased emotional egocentricity bias. In a female sample including both OC and NC women, Zhang et al. (2019) found no stress effect on pro-social decision-making, whereas men showed more pro-social behavior with higher cortisol stress responses. Both studies did not measure steroid hormone levels however, making a direct comparison with our findings difficult. There is also extensive literature about stress effects on empathy (Nitschke & Bartz, 2023) and some evidence for sex differences in these effects. For instance, Nitschke and colleagues (2022) found no stress effects on empathic accuracy in women, whereas men were more accurate after stress compared to before stress. Interestingly, this study also reported generally reduced empathic accuracy in OC users compared to NC women. Sex differences in stress effects on empathy seem to be context dependent however, as others found a stress-induced reduction in empathic accuracy in women compared to men (Crenshaw et al., 2019), when assessed in romantic couples. Otherwise, social cognition even increased in women after stress, but only in low cortisol responders (Smeets et al., 2009). In sum, these previous studies suggest interactive effects of sex/gender and stress on social cognition and empathy, but the direction of effects seems to depend on the experimental measure of social cognition and affect. It is important to emphasize that our paradigm measures automatic emotional action control and thus addresses very different affective and neural processes than paradigms of empathic accuracy or social cognition. Clearly, more research is needed on how steroid hormones and stress interact in their effects on various cognitive and affective processes and how this ultimately affects social behavior.

Remarkably, neither NC and OC women nor women taking androgenic and antiandrogenic OC differed significantly in their cortisol stress response. This is surprising as a blunted cortisol stress response has been often replicated in previous studies (Lewis et al., 2019; Tronson & Schuh, 2022). However, there are also other reports who did not find any group differences in this stress marker (Lovallo et al., 2019; Sharma et al., 2020). At least Lovallo and colleagues found the reduced cortisol stress response in OC users only when tested in the morning, but not in the afternoon (Lovallo et al., 2019). As we tested participants in the afternoon, this might explain the lack of group differences. Moreover, we instructed participants not to exercise, eat anything, drink coffee, take medications, or smoke for one hour before the appointment, following previous studies (Roelofs et al., 2005). The time window of one hour might have been too short (Hansen et al., 2012), which might have introduced variability within the groups that prevented us from detecting group differences.

Finally, we could not analyze cortisol levels in 20 women which reduced power to detect group differences. Since cortisol is only one of many stress markers, the groups might still have differed in other stress responses such as their subjective stress experience or other physiological stress markers. Unfortunately, we did not measure any other stress responses in our participants. Because we focused on the approach and avoidance behavior of OC users and the potential modulating effect of stress-induced cortisol changes, we had no specific research questions or hypotheses regarding the subjective experience of stress. Previous work on OC effects on subjective stress experience or other stress markers is inconclusive (Lewis et al., 2019), as some studies reported no OC related differences in subjective stress responses (Kirschbaum et al., 1999; Merz, 2017; Mordecai et al., 2017; Sharma et al., 2020) but others did not measure it. More research is clearly needed to understand the effects of OC usage on stress responses on a subjective and behavioral level.

### 4.4 Limitations

Like many other studies investigating the influence of oral contraceptives on the female organism, our study is also limited by the “survivor effect” (Kutner & Brown, 1972). This means that our study only comprised women who did not stop taking OC because of serious side effects, whereas those who did develop side effects abandoned the treatment early. Future studies should implement a longitudinal, within-subject design to avoid pre-selection of participants. To be able to directly relate behavioral effects in the experimental paradigms to potential affective side effects, one should also systematically assess emotional well-being and psychiatric symptoms. The cross-sectional study design also means that the effects are not uniquely attributable to OC use but may be influenced by other factors which caused women to decide for or against OC. Although the lack of group differences in personality and emotion regulation questionnaires speaks against this argument, we cannot rule out the possibility that there are influencing variables which differ between women taking oral contraceptives or not.

In our AAT paradigm, we only presented faces with positive and negative valence. It would be beneficial to add a neutral condition in future studies to clarify if the shortened reaction times of OC women are due to the affective component. Combining the experiment with fMRI measurements could also shed light on the neural mechanisms associated with these effects. In particular, the amygdala and aPFC could be examined more closely to understand the neural basis of the observed behavioral changes (Kaldewaij et al., 2017).

We did not measure steroid hormone levels in the participants, which limits the conclusions we can derive from our observations. Steroid hormone measurements would have allowed to go beyond a simple group comparison and to directly correlate behavioral effects with hormonal levels. This could have helped to clarify the relative contribution of estradiol and progesterone to the AAT effects (for comparison, see Strojny et al. (2021)). Moreover, hormone measurements could have supported the presumed group differences in steroid hormones. We decided to test participants on the 1^st^ to 3^rd^ day of the active pill phase and, in case of women not taking hormonal contraception, on the 8^th^ to 10^th^ day of their menstrual cycle. As the 8^th^ to 10^th^ day are not clearly early follicular phase anymore, this might have led to more variability within the NC group already, as in some estradiol might have been rising already again. Also, we did not track cycle length of the participants but only asked them to come to the lab at the specified day after menses. However, such variability should rather have led to smaller effects or null effects. Also with regard to the measurement time in the OC group, there is debate on when to best perform measurements to study OC effects (Gravelsins et al., 2023). Hormonal levels might vary depending on whether women are in their active or inactive phase, when in the inactive phase and, depending on the pharmacokinetics, when during the day (Gravelsins et al., 2023; Stanczyk et al., 2013). In our study, we decided to keep the measurement time within the active phase constant, and we did not observe differences between participants depending on pill intake. However, it should be noted that hormone measurements could have clarified the hormonal status of participants and the lack of hormone measurements presents a limitation of this study.

### 4.5 Conclusion

This work stands in line with a series of previous studies showing that oral contraceptives can influence cognitive and affective processes (Gravelsins et al., 2023; Lewis et al., 2019; Tronson & Schuh, 2022). Specifically, we showed that OC use is associated with a reduced avoidance tendency to social threat signals and an increased stress-induced approach tendency to positive social signals. This expands on other research demonstrating altered reactivity to particularly negative emotional stimuli in OC users (Lewis et al., 2019) and suggests that these differences are not limited to the perceptual level, but have motivational implications. In addition to recent work speaking for a negative OC effect on affective empathy and prosocial decision-making (Kimmig et al., 2022; Strojny et al., 2021), we show that women taking oral contraceptives differ already on automatic motivated behaviors to affective signals. Future studies will have to examine how these differences relate to neural mechanisms on the one hand and presumed affective side-effects of OCs on the other hand.

## CRediT author statement

**Jasmin Thurley:** Methodology, Software, Formal analysis, Investigation, Writing – Original Draft, Visualization **Macià Buades-Rotger:** Methodology, Writing – Review and Editing **Georg Serfling:** Methodology, Writing – Review and Editing **Thessa Howaldt:** Investigation, Writing – Review and Editing **Nicole Reisch:** Writing – Review and Editing **Ulrike Krämer:** Conceptualization, Methodology, Software, Resources, Writing – Review and Editing, Supervision, Project administration

## Acknowledgements

We thank Carla Bücker, Annika Kühle and Luisa Ackermann for help during data acquisition.

UMK and NR are supported by the German Research Foundation (DFG) as part of the CRC 1665.

