## Supplementary Tables for "Reduced threat avoidance but increased stress induced approach bias in women taking oral contraceptives"

### Supplementary Material

**Supplementary Table 1: Composition of oral contraceptives used by participants**

| | number (participants) |
|---|---|
| Estrogen | |
| Ethinylestradiol (0,02) | 17 |
| Ethinylestradiol (0,03) | 41 |
| Estradiolvalerat (1, 2, 3) | 4 |
| Estetrol-Monohydrat (14,2) | 3 |
| Progestin | |
| Chlormadinon acetat (2) | 6 |
| Desogestrel (0,15) | 1 |
| Dienogest (2, 3) | 4 |
| Dienogest (2) | 24 |
| Drospirenon | 4 |
| Levonorgestrel (0,1) | 17 |
| Levonorgestrel (0,125) | 2 |
| Levonorgestrel (0,15) | 7 |

Notes: Number of participants for each estrogen and progestin component. The numbers in brackets refer to the dose in mg.

**Supplementary Table 2: Summary of side effects reported by participants**

| | number (participants) |
|---|---|
| Affective | |
| Mood swings | 6 |
| Depressive symptoms | 4 |
| Irritability | 2 |
| Libido (reduced) | 3 |
| Physical | 6 |
| Reduced menstrual pain | 2 |

Notes: Summary of side effects reported by participants. Physical symptoms included tension in the chest, nausea, headache, digestive symptoms and pain during sexual intercourse.
